# AAV-DJ is Superior to AAV9 for Targeting Brain and Spinal Cord, and De-Targeting Liver Across Multiple Delivery Routes in Mice

**DOI:** 10.1101/2024.05.06.592798

**Authors:** Monika Chauhan, Audrey L. Daugherty, Fatemeh (Ellie) Khadir, Ozgun F. Duzenli, Alexandra Hoffman, Jennifer A. Tinklenberg, Peter B. Kang, George Aslanidi, Christina A. Pacak

## Abstract

Highly efficient adeno associated viruses (AAVs) targeting the central nervous system (CNS) are needed to deliver safe and effective therapies for inherited neurological disorders. The goal of this study was to compare the organ-specific transduction efficiencies of two AAV capsids across three different delivery routes. We compared AAV9-CBA-f*LucYFP* to AAV-DJ-CBA-f*LucYFP* using the following delivery routes in mice: intracerebroventricular (ICV) 1x10^12^ vg/kg, intrathecal (IT) 1x10^12^ vg/kg, and intravenous (IV) 1x10^13^ vg/kg body weight. Our evaluations revealed that following ICV and IT administrations, AAV-DJ demonstrated significantly increased vector genome (vg) uptake throughout the CNS as compared to AAV9. Through the IV route, AAV9 demonstrated significantly increased vg uptake in the CNS. However, significantly fewer vgs were detected in the off-target organs (kidney and liver) following administration of AAV-DJ using the IT and IV delivery routes as compared to AAV9. Distributions of vgs correlate well with transgene transcript levels, luciferase enzyme activities, and immunofluorescence detection of YFP. Overall, between the two vectors, AAV-DJ resulted in better targeting and expression in CNS tissues paired with de-targeting and reduced expression in liver and kidneys. Our findings support further examination of AAV-DJ as a gene therapy capsid for the treatment of neurological disorders.

## Introduction

More than 520 gene therapy trials currently listed on clinicaltrials.gov are underway in the United States to investigate a wide variety of gene delivery vehicles for both gene replacement and genome editing therapeutic strategies. The gene therapy field has made significant advancements over the last few decades as there are now Food and Drug Administration (FDA) approved therapies for multiple genetically inherited disorders that were once considered untreatable (1–9).

Adeno-associated virus (AAV) is one of the most efficient vehicles available to deliver healthy gene cDNA sequences into the cells of affected individuals due to its ability to persist for long periods of time as an episome within the nuclei of cells, its relatively non-pathogenic nature, and its ability to infect dividing and non-dividing cells *in vivo* (10,11). Despite these advantages, one key challenge related to the use of AAV includes the high doses currently required to achieve a sufficient therapeutic effect. High doses of intravenously administered AAV can elicit responses in both the innate and adaptive immune systems (12–14). There are both vector-dependent and host-dependent factors that influence the immunogenicity of AAV capsids and subsequent severity of immune responses (14,15). Major obstacles related to immune responses include the presence of host neutralizing antibodies against AAV (16) and the activation of CD4^+^ and CD8^+^ T cells in response to degradation of either capsid or transgene-encoded proteins by the proteosome (17). These inflammatory responses can be further amplified by activation of the alternative complement pathway through direct interactions between complement component C3 and AAV capsid proteins (12). Efforts to reduce the therapeutic dose to help increase safety, reduce immune responses, and improve the therapeutic efficacy of gene therapies promise to yield significant clinical benefits. One approach to improve AAV-based therapeutics is to develop capsid serotypes that are optimal for transducing desired organs and cell types, while simultaneously avoiding off-target organs.

Our interest in the development of AAV-mediated gene-based therapies for rare neurological disorders led us to explore novel capsids for targeting specific regions and cells within the brain and spinal cord. Genetic disorders associated with neurological pathologies are particularly difficult to treat due to challenges associated with penetration of the highly protective blood-brain barrier (BBB) and central nervous system (CNS) complexities that can result in limited access to the brain’s deep structures. AAV capsid motifs that are unique for each serotype are critical for binding to host cells and represent a key step in the specificity of AAV-mediated gene delivery. We thus compared the AAV9 capsid currently used in the FDA approved gene therapy for spinal muscular atrophy (SMA) (onasemnogene abeparvovec-xioi) to the AAV-DJ capsid that was recently demonstrated to be superior for the broad transduction of neurological tissue in an extensive non-human primate study (18).

Another important consideration is the delivery route for gene therapy. Thus, we also compared the efficiency of AAV9 and AAV-DJ capsids across three different delivery routes with respect to their ability to target the CNS and simultaneously de-target the liver and kidney. We are particularly keen to avoid the liver as AAVs that are delivered systemically tend to be sequestered in the liver (19,20), concurrently reducing transduction in target organs and potentially leading to liver toxicity (21,22), which may in some cases be fatal (23). Our study will help optimize capsid selection and delivery route for CNS gene therapies.

## Materials and Methods

### AAV Production and Purification

Both AAV9 and AAV-DJ vectors were packaged in a single-strain expression cassette with chicken β-actin (CBA) promoter-driven fusion of firefly luciferase (Luc), and yellow fluorescent protein (YFP) (24,25). Recombinant AAV vectors were produced using the triple transfection method and purified as described previously (26,27). Briefly, HEK293 cells were co-transfected with three plasmids encoding 1) Rep/Cap, 2) ITRs-CBA-*fLuc-YFP*, and 3) Ad helper genes using polyethyleneimine (#23966-1, Polysciences, USA). Cells were harvested 72 hrs post transfection, subjected to freeze-thaw cycle, and treated with Benzonase (#E8263, Sigma-Aldrich, USA) at 37°C for 1 hr. The suspension was purified by Iodixanol (#D1556, Sigma-Aldrich, USA) gradient ultracentrifugation followed by ion exchange chromatography using HiTrap^®^ Q HP (Cytiva, USA) with Bis-Tris propane-MgSO_4_ buffer. Subsequently, the vector preparation was concentrated into 10 mM Tris – 100 mM sodium citrate buffer (pH 7) using centrifugal spin concentrators (#AP2015010, Orbital Biosciences, USA).

### Quantitative PCR analysis for the determination of AAV titers

For vector genome (vg) titer determination, DNA contaminants were removed by TURBO DNase (#REF4022G, Thermo Fisher Scientific, USA) digestion. AAV genomes were then released from AAV capsids by Proteinase K (#AM2546, Invitrogen, USA) digestion. Subsequently, the viral DNA was cleaned using DNA Clean & Concentrator^TM^ 25 (#11-305C, Genesee Scientific, USA). Vector titers were quantified by qPCR with a TBGreen^®^ Advantage (#S4748, Takara Bio, USA), using the following primer pair specific to the CBA promoter region within the viral cassette: forward primer 5’-TCCCATAGTAACGCCAATAGG -3’ and reverse primer 5’- CTTGGCATATGATACACTTGATG -3’(26,27).

### Animals

All husbandry and procedural use of animals was approved by the Institutional Animal Care and Use Committee (IACUC) at the University of Minnesota (UMN). All the FVB/NJ (stock# 001800) mice (5 weeks old) were ordered from Jackson Laboratory. The mice were maintained at 25°C and 50% humidity in the Research Animal Resources facility at UMN. Animals were maintained in 14/10 h light/dark cycles and fed chow and water ad libitum. All animals used in the study were acclimatized in the vivarium and handled for one week prior to initiation of the study.

### Routes of AAV delivery

Equal numbers of males and females were used for each delivery route and capsid. In total, for every route, 10 (5 males and 5 females) 6-week-old mice were injected with AAV9-CBA- f*Luc/YFP* and 10 (5 males and 5 females) 6-week-old mice were injected with AAV-DJ-CBA- f*Luc/YFP*.

### Jugular Vein Injection (IV)

IV administration was performed as previously described (28,29). Briefly, mice were anesthetized with inhaled 1% to 3% isoflurane (Dechra, USA). A small incision (∼0.5 cm) was made parallel to the midline at the lower third of the right anterior neck, exposing the right external jugular vein. A 28-gauge insulin needle was bent on the sterile surface and used to slowly inject 150 µL of AAV-CBA-f*Luc/YFP* or vehicle (1x PBS) into the vein. Gentle pressure was applied on the skin to achieve hemostasis and the incision was closed with the tissue adhesive (Vetbond 1469, 3M animal care products, USA). 1x10^13^ vg/kg body weight per mouse was injected in a total volume of 150 µL. Meloxicam (OstiLox, VETone, UK) at 0.5mg/kg was injected subcutaneously as an analgesic for 72 hrs post-surgery. Animals were monitored carefully post-surgery according to IACUC instructions.

### Intrathecal Injection (IT)

For intrathecal injections, mice were anesthetized as mentioned above. Hair was removed from a small area on the lower back and scrubbed. A Hamilton syringe (26 gauge) was used to deliver the AAV (1x10^12^ vg/kg body weight per mouse) directly by lumbar puncture (between the fifth lumbar (L5) and the sixth lumbar (L6) vertebrae) in a total volume of 10 µL (30–33). Male and female cohorts were injected for both capsids separately as mentioned above.

### Intracerebroventricular Injection (ICV)

For bilateral ICV injections, mice were anesthetized, hair was removed from the area between the ears, the head was fixed into the stereotaxic apparatus and scrubbed. A small incision was made, a T-shaped subcutaneous junction (bregma) was identified, a 26 G Hamilton needle attached to the apparatus was adjusted to that point, and subsequently the apparatus values for dorsoventral axis were set to zero. The needle was used to direct to the lateral ventricles: -0.5 mm in the anteroposterior axis, ±1 mm in the mediolateral axis, and -2.3 mm in the dorsoventral axis (∼2 months old mouse coordinated according to Allen Brain Atlas). Once the needle was placed inside the ventricle, 5 µL of AAV-CBA-f*Luc/YFP* or vehicle (1x PBS) was injected slowly into the ventricle with a flow rate of 1 µL per minute to enable further diffusion (34). Before slow withdrawal of the needle, there was a 2 min wait time to ensure complete diffusion of the injected fluid and no back-flow of the delivered virus. Similarly, the other hemisphere was injected. The total injected AAV dose was 1x10^12^ vg/kg body weight per mouse. The incision was closed with tissue adhesive (3M Vetbond, 3M animal care products, USA), and meloxicam (OstiLox, VETone, UK) at 0.5mg/kg was injected subcutaneously as an analgesic for 72 hrs post-surgery. Animals were monitored carefully post-surgery according to IACUC instructions.

### *In Vivo* Live Imaging

Two weeks following AAV administration, *in vivo* imaging was performed to monitor luciferase expression in live animals. The mice were administered with D-luciferin (#770504, IVISbrite D-Luciferin, RediJect, PerkinElmer, USA) through intra-peritoneal (IP) injections at a dose of 150 mg/kg of body weight. After 15 min, mice were subjected to anesthesia (isoflurane inhalation) followed by bioluminescence imaging analysis using an *in vivo* optical imaging system (IVIS100 IVIS® Imaging System, PerkinElmer, USA). Raw images containing raw data were then analyzed in M3Vision software (Living Image Software, PerkinElmer), using the freehand tool to obtain total luciferase signals from each organ. Data were exported in photons (ph)/s/cm^2^/steradian unit and displayed as a pseudo-color overlay onto the animal image, using a rainbow color scale.

### AAV vector genome (vg) determination

AAV vg copy numbers from various mouse tissues were determined as previously described (28,29) by qPCR after extraction of total genomic DNA (gDNA) using DNeasy Blood and Tissue kit (#69506, Qiagen, Germany). Tissues were mechanically homogenized using a bead basher (BenchMark) with a lysis buffer kit. gDNA was isolated according to manufacturer instructions. Vg content in each tissue was determined using 100 ng total DNA using the qPCR method on the Applied Biosystems QuantStudio 3 thermal cycler and a Taqman probe specific for the luciferase reporter sequence (Assay ID: Mr03987587_mr, Thermo). Luciferase plasmid DNA dilutions with known copy number were used to create a standard curve to extrapolate absolute copy numbers in the tissue sample gDNA (35,36).

### Transcript analysis

Total RNA was isolated from mouse tissues using the Quick-RNA MiniPrep kit (#R1055, Zymo Research, USA) according to manufacturer’s instructions. The RNA kit includes a gDNA removal step. 300 ng of total gDNA-free-RNA was used for cDNA synthesis using the High-Capacity RNA-to-cDNA Kit (#4388950, Applied Biosystems, USA). This cDNA product served as the template for qPCR with luciferase probes (Assay ID: Mr03987587_mr, Thermo, USA) and the Applied Biosystems QuantStudio 3 thermal cycler. Ribosomal RNA 18s amplification (Assay ID: Mm03928990_g1, Thermo, USA) on the same cDNA was used as an internal control. Fold change in luciferase transcripts was calculated using the ΔΔC_T_ method.

### Luciferase Assay

The frozen tissues (∼25 mg) were homogenized in 400 µL of 1x reporter lysis buffer (#E1501 kit, Promega), and the lysates underwent 3 freeze thaw cycles (-80^°^C to 37°C). The samples were centrifuged for 3 min. at 10,000 x g and supernatants were collected for the assay. In a 96 well plate (white bottom) 20 µL lysate was mixed with 50 µL Luciferase Assay Reagent (#E1501 kit, Promega) (as per the manufacturer’s instructions). The samples were run in triplicates and luminescence was measured with a 10,000 ms integration time using the SpectraMax i3x plate reader (Molecular Devices).

### Immunohistochemistry

Animals used for immunohistochemistry studies were euthanized by perfusion fixation as previously described (37,38). Mice were deeply anaesthetized with isoflurane and perfused with sterile 1x PBS (pH 7.4) followed by a fixative (4% paraformaldehyde (PFA) in 1x PBS). Tissues were removed post fixation and placed in 4% PFA for 12 h overnight. The next day, the tissues were dehydrated and embedded into paraffin blocks. Sections were cut at 7 µm thickness and mounted onto pre-charged slides (#12-550-15 Superfrost Plus slides, Fisher Scientific, USA). For immuno-fluorescence (IF), tissue sections were rehydrated in 100% xylene, alcohol gradient (100%, 70%, 50%) and then water. Sections were permeabilized for 30 min at room temperature in 1x PBS containing 0.3% Triton X-100 and then incubated for 1 h in blocking solution (1x PBS containing 3% BSA, 2% normal donkey serum, and 0.15% Triton X-100). The same blocking reagent was used to prepare primary antibody dilutions which were incubated on samples overnight at 4°C. The primary antibodies used include: mouse monoclonal anti-GFP/YFP, 1:50 (#A11121, Invitrogen); specificity confirmed by lack of labeling in the absence of AAV-CBA- f*Luc/YFP* treatment); rabbit anti-NeuN, 1:100 (#26975-I-AP, Proteintech); rabbit anti-GFAP, 1:100 (#NB300-141, Novus Biologicals), exclusive staining of astrocytes; rabbit anti-OLIG2 (#13999-1-AP, Proteintech); rabbit anti-doublecortin (DCX), 1:100 (#4604, Cell Signaling). After a 5 min 1x PBS rinse, sections were incubated for 1 h at room temperature with appropriate combinations of Alexa Fluor (1:500, AF568 goat anti-rabbit #A11036 and AF488 goat anti-mouse #A11029, Invitrogen) conjugated secondary antibodies (Life Technologies, Invitrogen). Sections were rinsed again in 1x PBS. Following the final rinses, sections were cover-slipped using a mounting solution containing DAPI (#P36962, ProLong Diamond Antifade Mountant, Invitrogen). Tissues from 1x PBS sham-injected mice were processed in parallel with the AAV-CBA-f*Luc/YFP* treated tissues and were included in all immunohistochemical experiments to control for non-specific staining of the YFP antibody.

### Microscopy and Image Analysis

Images were collected on a Leica DM5500 B epi-florescent microscope with LAS-X software. For comparisons of tissues from ICV, IT, and IV injected mice, images were collected using the same microscope setting with a few exceptions where adjustments were necessary to allow detection of labeling in IT treated tissues or avoid saturation in ICV and IV treated tissues. Similarly, image adjustments for contrast, brightness, and color were performed in parallel for all delivery routes and treated tissues. For each mouse tissue, 5 non-overlapping images were taken across 3 tissue sections, which were evaluated by a trained, blinded observer. Different cell types in the brain were outlined based on visualization with cell-type specific antibodies, and only cells with a visible nucleus were counted. Total number of YFP positive cell were counted using the cell counter plugin in Image J software. The percentage of YFP positive cells were calculated from the total number of each cell type.

### Statistical analysis

The data are expressed as mean or percentage ± standard error of the mean (SEM) from at least three animals unless indicated otherwise. For *in vivo* mouse imaging experiments, 10 mice were analyzed per treatment. For vgs, RNA, and luciferase assays, tissues from 6 mice were used in each group. For all IF studies, tissues from three different animals were analyzed. All the bar graph results are shown as mean ± SEM. No data were excluded from analyses. For determining the cell type counting with immunofluorescence assay images, the study investigators A.L.D and A.H. were blinded to treatment groups using codes generated by an independent investigator (M.C.).

Five male and five female mice were used for each AAV delivery cohort. Comparisons were made using one-way analysis of variance (ANOVA) followed by a Bonferroni post-hoc test or by a two-tailed student t-test unless otherwise indicated. In figure legends, n indicates the number of animals used in the study (biological replicates). P values were calculated by the indicated statistical tests using GraphPad Prism (v.10.4.0) software.

## Results

For all *in vivo* studies to assess the biodistribution of AAV9 and AAV-DJ, we used a ubiquitous Chicken β-actin (CBA) promoter to drive the expression of a firefly luciferase (Luc) fused to a yellow fluorescent protein (YFP). This combination of promoter and transgene allowed for a versatile and highly sensitive evaluation of CNS expression for comparisons of each capsid using the intravenous (IV), intrathecal (IT), and intracerebroventricular (ICV) delivery routes in healthy wild-type FVB/NJ mice. For the ICV and IT routes, each mouse received 1x10^12^ vg/kg of either AAV9-CBA-f*Luc/YFP*, AAV-DJ-CBA-f*Luc/YFP,* or an equivalent volume of 1x PBS (for injection controls). For the IV route, each mouse received 1x10^13^ vg/kg of either AAV9-CBA- f*Luc/YFP*, AAV-DJ-CBA-f*Luc/YFP,* or an equivalent volume of 1x PBS (**Figure 1A**). No difference between males and females was observed. Mice from all cohorts were imaged for detection of luciferase mediated bioluminescence at three, seven, and nine weeks post-AAV administration (**Figure 1B**).

**Figure 1.**
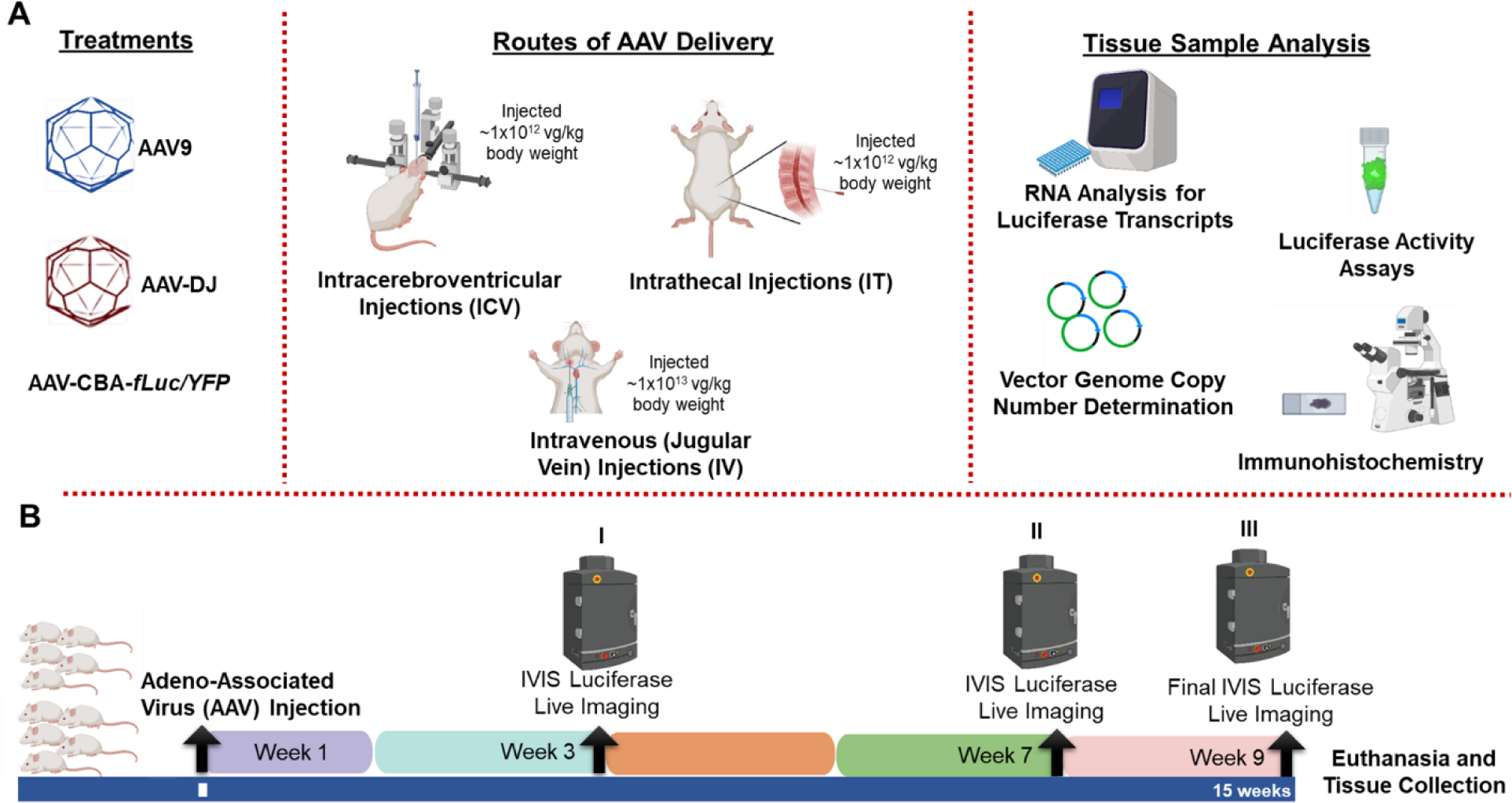
Experimental overview of capsid and delivery route comparisons. (A) WT FVBN/J mice at ∼6 weeks of age were injected via intracerebroventricular (ICV), intrathecal (IT), or jugular vein (IV) injections with AAV9 or AAV-DJ to deliver a luciferase - yellow fluorescent protein (YFP) fusion protein. (B) Mice were evaluated throughout the study through non-invasive luciferase live imaging at the end of weeks 3 (I), 7 (II) and 9 (III). Nine weeks after AAV delivery, six mice (3 males and 3 females) were euthanized for DNA, RNA, and protein analysis and four mice (2 males and 2 females) for histological studies.

Bioluminescence was detected from all AAV9 and AAV-DJ treated mice throughout every imaging session. As our goal was to primarily target the CNS, head/body radiance ratios were determined to provide relative biodistribution of expression information. Mice from both the ICV and IT AAV-DJ cohorts displayed a significant increase in relative head/body ratios at nine weeks as compared to those treated with AAV9 using the same delivery routes (**Figure 2A-D, Supp. Figure 1**). In comparison, mice from the IV delivery route that were administered AAV9 displayed significantly higher head/body ratios at nine weeks as compared to those in the AAV-DJ cohort (**Figure 2F**).

**Figure 2.**
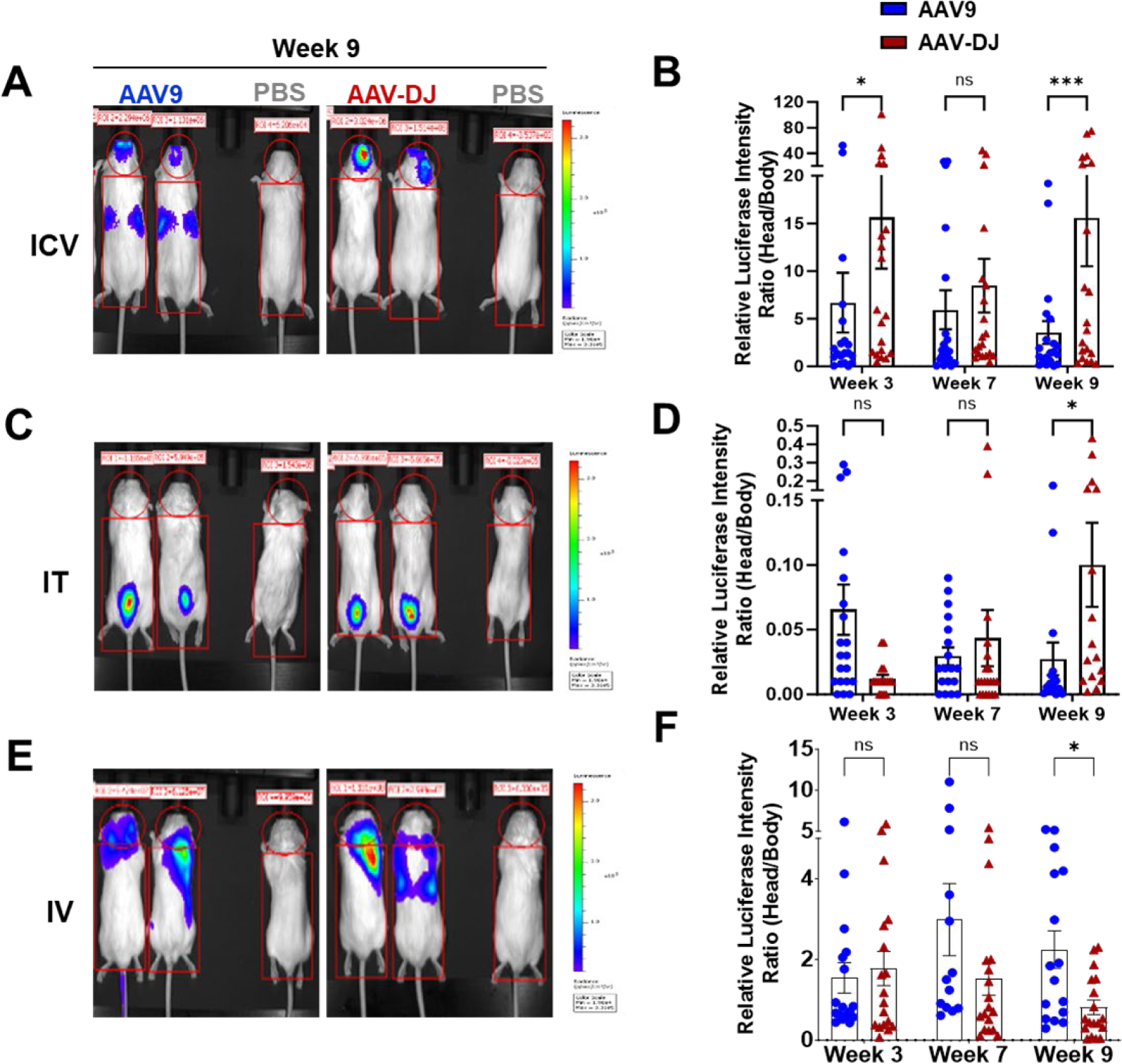
Visualization and quantification of differential Luciferase expression patterns using IVIS whole-body live bioluminescent imaging in WT FVBN/J mice. 1×10^12^ vg/kg (ICV or IT) or 1×10^13^ vg/kg (IV) of mouse body weight of AAV (AAV-CBA-*fLuc/YFP)* encoding luciferase gene was delivered into 6 weeks old mice via (A) ICV, (C) IT, or (E) IV. Luciferase expression levels were detected by injecting luciferin substrate 15 min prior to bioluminescence imaging at 3, 7, and 9 weeks after vector administrations. Representative images of two mice from AAV9 and AAV-DJ groups for Week 9 post AAV delivery are shown. Mouse 1 from the right side of each image is the No virus control. Luciferase intensities from both the dorsal and ventral sides of each animal were obtained from the bioluminescent images and head to body ratios were calculated and presented in the graphs (B) (ICV), (D) (IT) and (F) (IV). The data are shown as mean values ± SEM (N=10 mice per group; *p < 0.05, ***p < 0.001, ns- not significant based upon unpaired two-tailed t-tests).

To quantify both the efficiency of gene transfer to various regions of the CNS and de-targeting of liver and kidney, full necropsies were performed on six mice from each cohort at nine weeks post-AAV administration. gDNA was isolated and AAV vector genome (vg) biodistribution was determined across the following brain regions: hippocampus, hypothalamus, cerebellum, and cortex, as well as spinal cord, liver, and kidney (**Figure 3**). Using the IT and ICV routes, vg levels in the AAV-DJ cohorts were significantly higher than those from AAV9 cohorts in the hippocampus, hypothalamus, cerebellum, cortex, and spinal cord (**Figure 3A-E**). In contrast, IT administration of AAV9 resulted in significantly higher levels of vgs in the off-target liver and kidney while ICV administration showed no differences between the two capsids (**Figure 3F, G**). Using the IV delivery route, mice in the AAV9 cohort displayed significantly higher vg levels in the hypothalamus, cerebellum, and cortex as well as in the off-target organs liver and kidney as compared to AAV-DJ. IV delivery with the AAV-DJ capsid resulted in significantly higher vg levels in the hippocampus and spinal cord as compared to AAV9.

**Figure 3.**
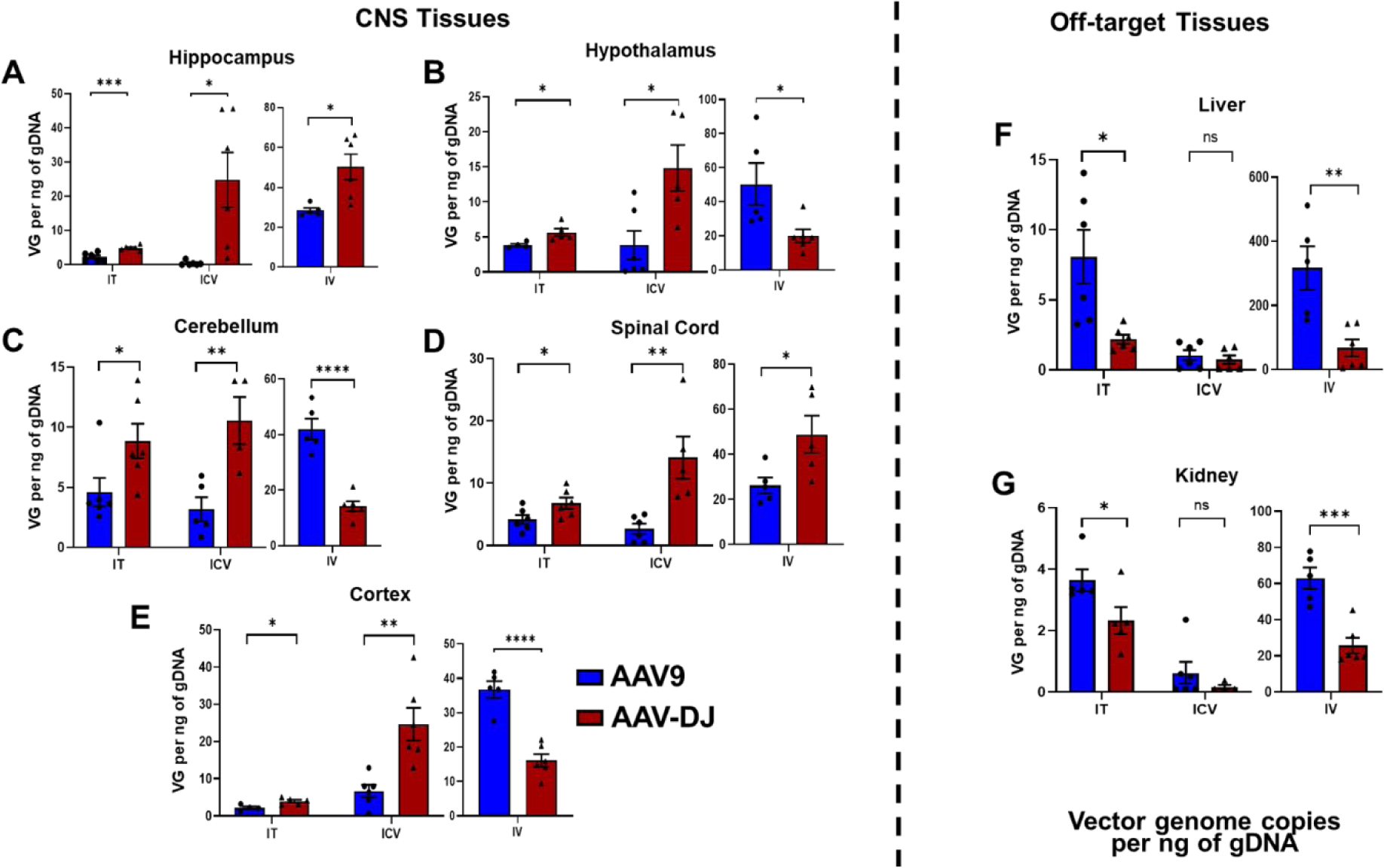
AAV vector genome (VG) estimations in tissues of interest. gDNA was isolated and luciferase gene vector genome numbers were calculated per nanograms of gDNA for (A-E) neurological tissues (hippocampus, hypothalamus, cortex, cerebellum, spinal cord) and off-target tissues (F) (liver) and (G) kidney. Data are presented as mean ± SEM of the values from six mice; *p < 0.05, **p < 0.01, ***p < 0.001, ns- not significant (based upon unpaired two-tailed t-tests).

To further examine the efficiency of gene delivery by AAV-DJ and AAV9, luciferase gene transcription levels were assessed in the CNS and off-target organs. Total RNA was isolated from each tissue and luciferase transcription levels were assessed by quantitative PCR. The hippocampus, cerebellum, cortex, and spinal cord from the IT groups demonstrated significantly higher transcription levels following delivery with AAV-DJ whereas the hypothalamus, liver, and kidney had higher levels following delivery with AAV9 (**Figure 4 A-G**). The hippocampus, hypothalamus, cerebellum, spinal cord, and cortex from the ICV group all demonstrated significantly higher transcription levels following delivery with AAV-DJ whereas the liver and kidney showed no significant differences in expression levels between AAV-DJ and AAV9. Every neurological tissue analyzed except the cerebellum showed significantly higher expression following the use of IV administered AAV9 (**Figure 4A-E**). However, both off-target organs (liver and kidney) also showed significantly higher expression levels with AAV9 (**Figure 4F, G**).

**Figure 4.**
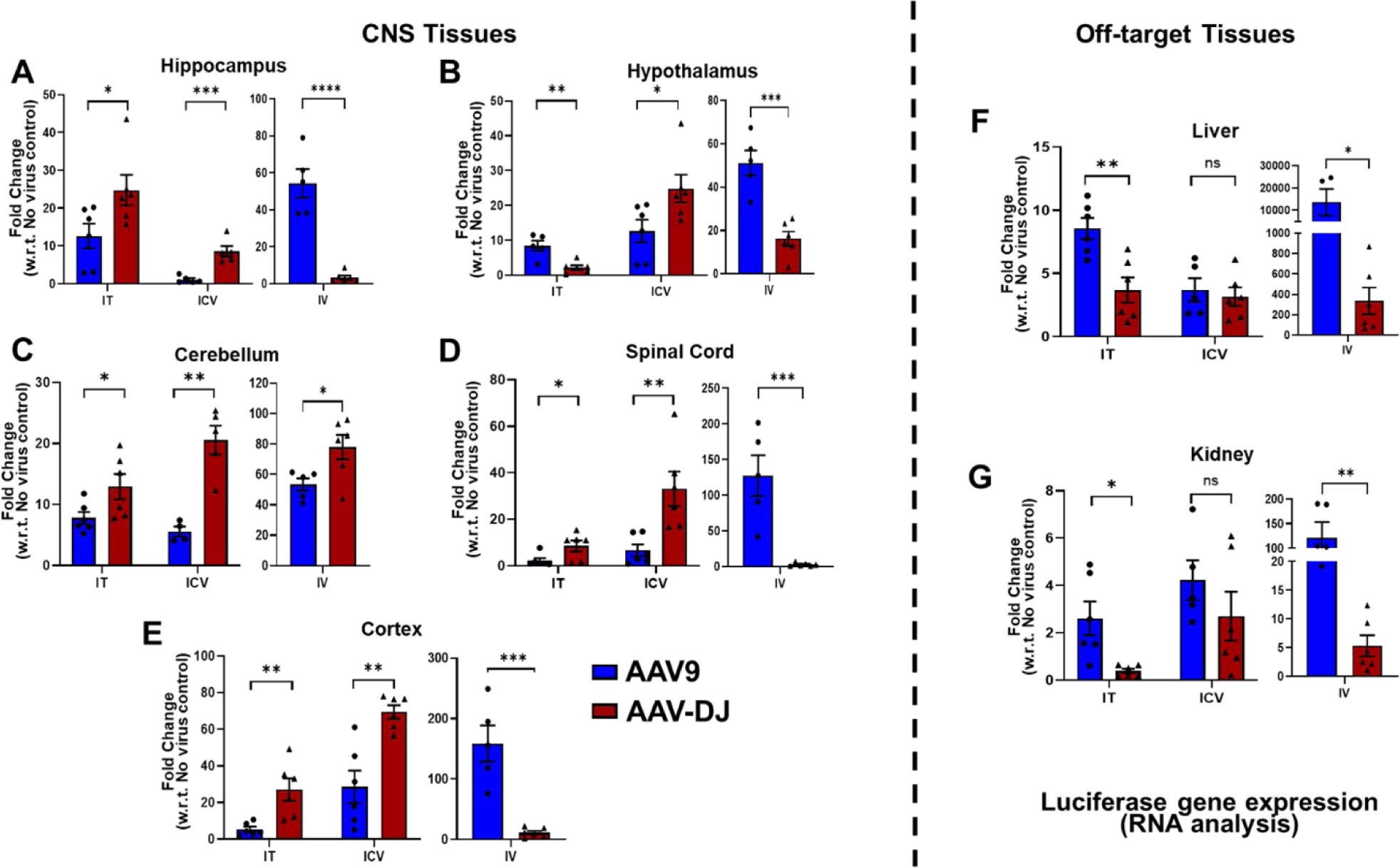
Luciferase transgene transcript levels in tissues of interest. Total RNA was isolated, converted to cDNA, and fold changes in luciferase mRNA transcript expression levels were calculated as compared to no virus controls for (A-E) neurological tissues (hippocampus, hypothalamus, cortex, cerebellum, spinal cord) and off-target tissues (F) (liver) and (G) kidney. Data are presented as mean ± SEM of the values from six mice; *p < 0.05, **p < 0.01, ***p < 0.001, ns- not significant (based upon unpaired two-tailed t-tests).

Luciferase activity assays were used to quantify relative luciferase protein levels among organs and cohorts. Relative light units (RLUs) were normalized to the amount of total protein for each respective sample. The hippocampus, cerebellum, cortex, and spinal cord from the IT groups demonstrated significantly higher luciferase activity levels following delivery with AAV-DJ whereas the hypothalamus, liver, and kidney had higher levels following delivery with AAV9 (**Figure 5A-G**). The hippocampus, hypothalamus, cerebellum, cortex, and spinal cord from the ICV group all demonstrated significantly higher luciferase activity levels following delivery with AAV-DJ whereas the liver showed higher levels with AAV9 and the kidney showed no significant differences in expression levels between AAV-DJ and AAV9 (**Figure 5A-G**). Every neurological tissue analyzed except the spinal cord showed significantly higher activity levels following the use of IV administered AAV9 (**Figure 5A-E**). Both off-target organs (liver and kidney) also showed significantly higher levels with AAV9 (**Figure 5F, G**).

**Figure 5.**
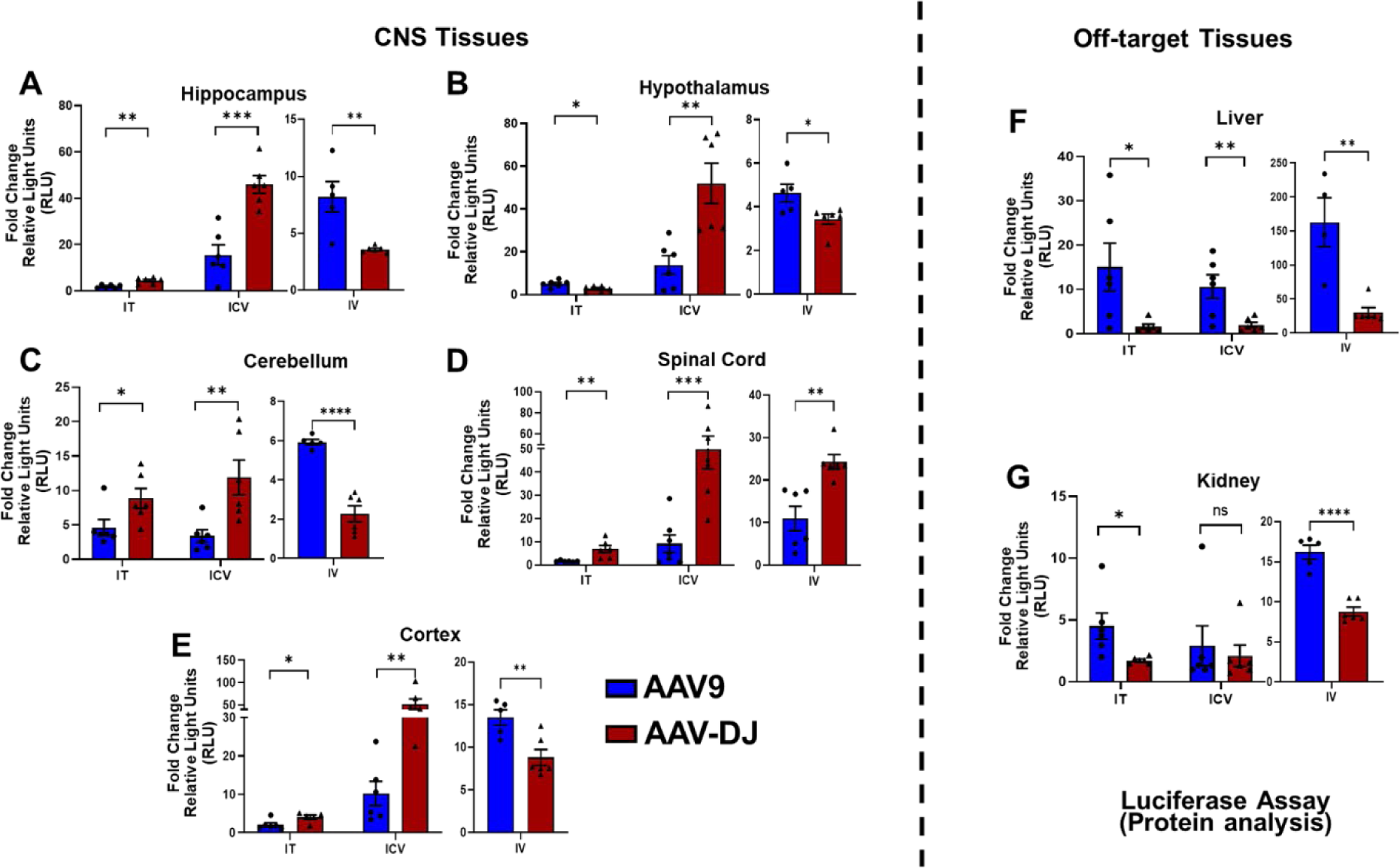
Luciferase activity assays in tissues of interest. Relative luciferase activities were measured and normalized to total protein concentrations and presented as fold changes in RLUs were calculated as compared to no virus controls for (A-E) neurological tissues (hippocampus, hypothalamus, cortex, cerebellum, spinal cord) and off-target tissues (F) (liver) and (G) kidney. Data are presented as mean ± SEM of the values from six mice; *p < 0.05, **p < 0.01, ***p < 0.001, ****p < 0.0001, ns- not significant (based upon unpaired two-tailed t-tests).

Immunofluorescence (IF) staining was performed to identify differences in the ability of AAV9 and AAV-DJ to transduce specific neurological cell types. Co-immunostaining to detect YFP transgene expression and molecular markers of neurons (neuronal nuclei, NeuN), oligodendrocytes (olig2), and astrocytes (glial fibrillary acidic protein, GFAP) was performed on fixed brain slices from mice representing each cohort. To facilitate quantitative evaluations of gene transfer efficacies between cohorts, the numbers of YFP/NeuN, YFP/olig2, and YFP/GFAP double-positive cells among the total number of neurons, oligodendrocytes, or astrocytes (respectively) were determined following random image sampling of coronal mouse brain sections (**Supp. Figure 3, 4 and 5**).

Both the AAV9-CBA-f*Luc/YFP* and AAV-DJ-CBA-f*Luc/YFP* vectors yielded a similar general distribution of expression in neurons across brain regions (**Supp. Figure 3** and **Figure 6**). Following ICV administrations, YFP expression was detected in 5.32% ± 3.98% (AAV9) and 11.77% ± 8.93% (AAV-DJ) of NeuN+ cells (**Figure 6A**). Following IT administrations, YFP expression was detected in 5.07% ± 2.83% (AAV9) and 6.50% ± 5.60% (AAV-DJ) of NeuN+ cells (**Figure 6B**). Following IV administrations, YFP expression was detected in 8.43% ± 3.21% (AAV9) and 5.46% ± 2.16% (AAV-DJ) of NeuN+ cells (**Figure 6C**).

**Figure 6.**
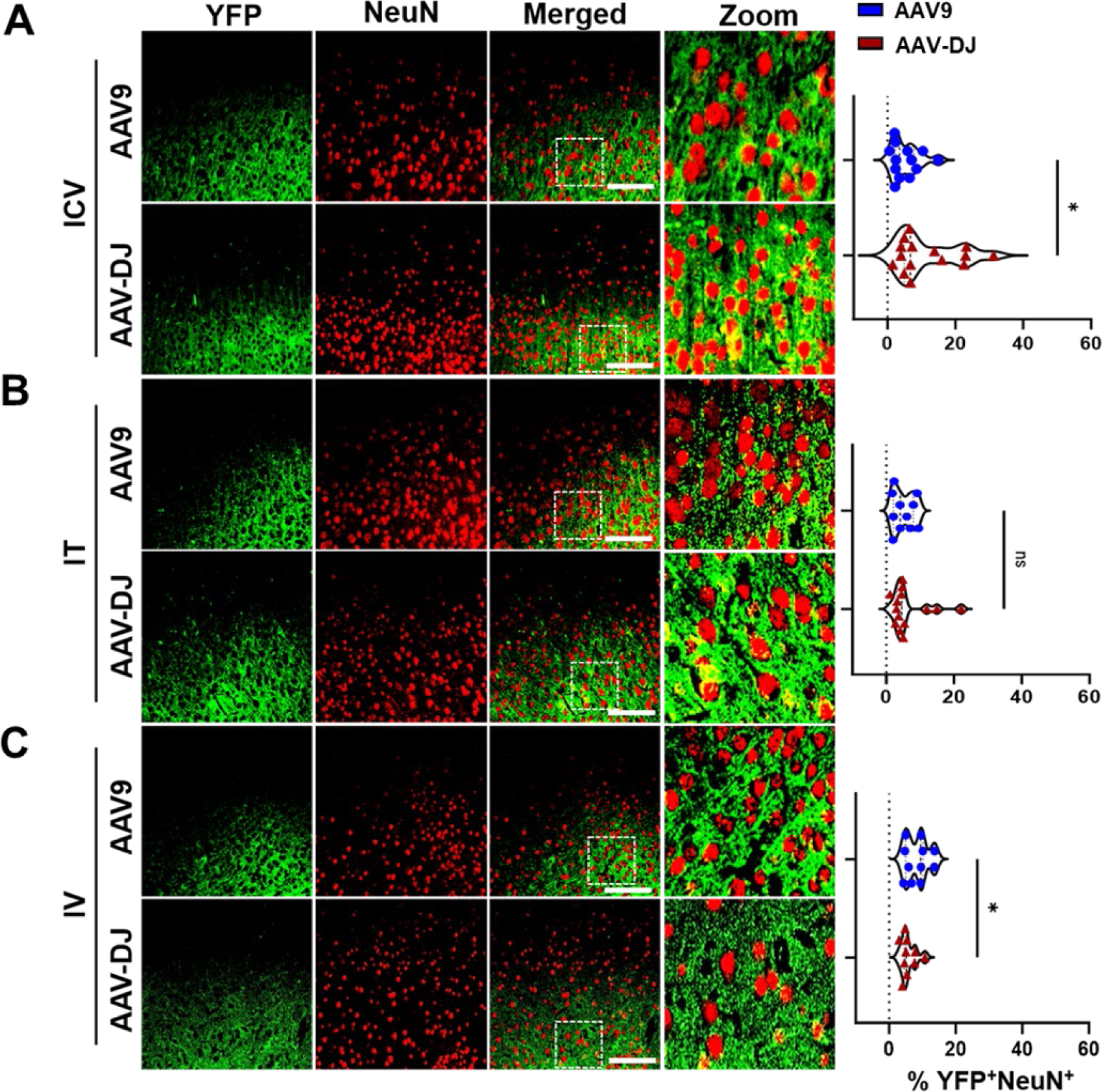
Relative transduction of neurons between cohorts. YFP transgene expression was compared to a neuronal cell marker (NeuN) and the co-localization was quantified across cohorts through blinded analysis. Representative immunofluorescence images from each cohort (A) ICV, (B) IT, and (C) IV are provided. For each mouse tissue, 5 non-overlapping images were acquired from across 3 tissue sections, and evaluated by a trained, blinded observer. The percentages of YFP^+^/NueN^+^ cells are presented in the right-hand panels. Scale bars = 100 µm. The white box indicates the zoomed-in portion of the image that is displayed to the right of each row of images.

Both the AAV9-CBA-f*Luc/YFP* and AAV-DJ-CBA-f*Luc/YFP* vectors yielded a similar general distribution of expression in oligodendrocytes across brain regions (**Supp. Figure 4** and **Figure 7**). Following ICV administrations, YFP expression was detected in 12.19% ± 7.18 % (AAV9) and 19.61% ± 7.88 % (AAV-DJ) of olig2^+^ cells (**Figure 7A**). Following IT administrations, YFP expression was detected in 16.45% ± 9.01% (AAV9) and 14.32% ± 5.48% (AAV-DJ) of olig2+ cells (**Figure 7B**). Following IV administrations, YFP expression was detected in and 16.82% ± 10.49% (AAV9) and 9.92% ± 5.13% (AAV-DJ) of olig2^+^ cells (**Figure 7C**).

**Figure 7.**
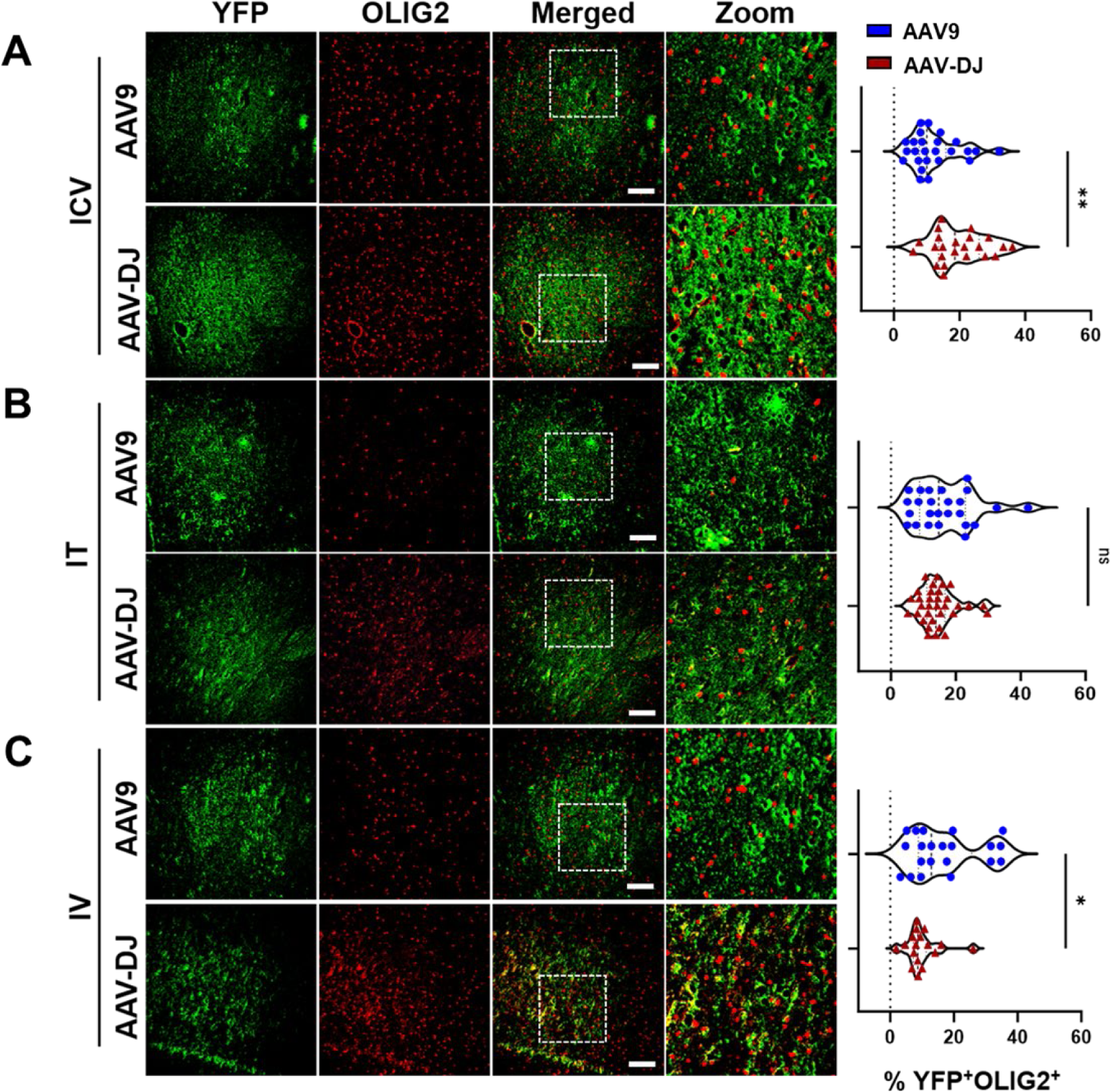
Relative transduction of oligodendrocytes between cohorts. YFP transgene expression was compared to an oligodendrocyte cell marker (OLIG2) and the co-localization was quantified across cohorts through blinded analysis. Representative immunofluorescence images from each cohort (A) ICV, (B) IT, and (C) IV are provided. For each mouse tissue, 5 non-overlapping images were acquired from across 3 tissue sections, and evaluated by a trained, blinded observer. The percentages of YFP^+^/OLIG2^+^ cells are presented in the right-hand panels. Scale bars = 100 µm. The white box indicates the zoomed-in portion of the image that is displayed to the right of each row of images.

Both the AAV9-CBA-f*Luc/YFP* and AAV-DJ-CBA-f*Luc/YFP* vectors yielded a similar general distribution of expression in astrocytes across brain regions (**Supp. Figure 5** and **Figure 8**). Following ICV administrations, YFP expression was detected in 14.15% ± 4.28% (AAV9) and 17.49% ± 4.97% (AAV-DJ) of GFAP+ cells (**Figure 8A**). Following IT administrations, YFP expression was detected in 14.46% ± 6.13% (AAV9) and 15.89% ± 5.84% (AAV-DJ) of GFAP+ cells (**Figure 8B**). Following IV administrations, YFP expression was detected in 12.28% ± 5.40% (AAV9) and 16.92% ± 6.93% (AAV-DJ) of GFAP+ cells (**Figure 8C**).

**Figure 8.**
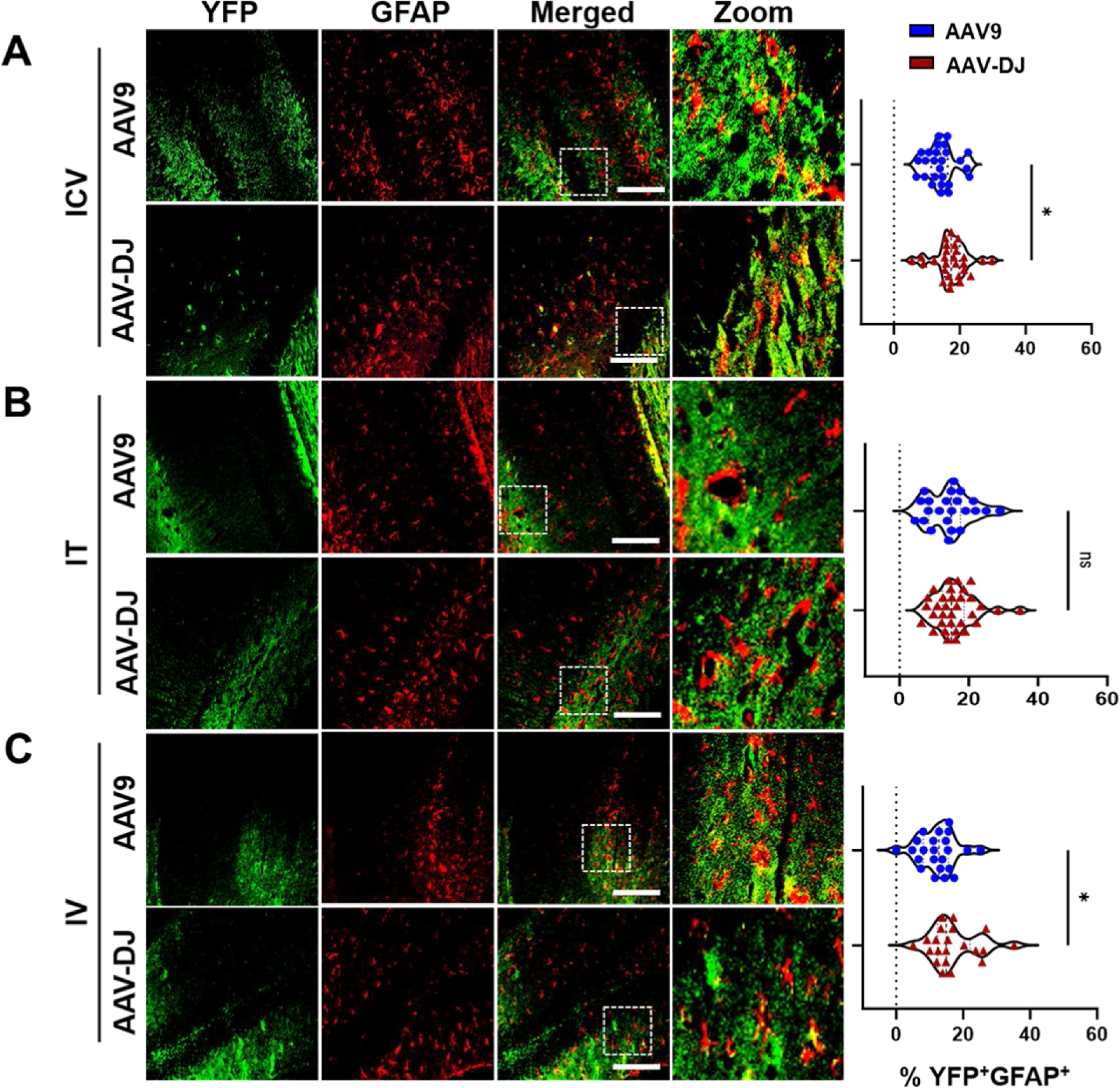
Relative transduction of neurons between cohorts. YFP transgene expression was compared to an astrocyte cell marker (GFAP) and the co-localization was quantified across cohorts through blinded analysis. Representative immunofluorescence images from each cohort (A) ICV, (B) IT, and (C) IV are provided. For each mouse tissue, 5 non-overlapping images were acquired from across 3 tissue sections, and evaluated by a trained, blinded observer. The percentages of YFP^+^/GFAP^+^ cells are presented in the right-hand panels. Scale bars = 100 µm. The white box indicates the zoomed-in portion of the image that is displayed to the right of each row of images.

Based upon these data, the AAV-DJ capsid transduces significantly more neurons, astrocytes, and oligodendrocytes than the AAV9 capsid when using the ICV administration route. No significant differences were observed between the ability of the two capsids to transduce neurons, astrocytes, or oligodendrocytes following IT administrations. In contrast, the IV administration data showed more variation with the AAV9 capsid transducing significantly more neurons, AAV-DJ transducing significantly more astrocytes, and no significant difference observed in the ability of these capsids to transduce oligodendrocytes.

To generate an overall assessment of our findings, we generated a heat map that included a score for each experimental result based on the goal of optimizing transduction of neurological tissue and de-targeting the liver and kidney (**Figure 9**). This heat map indicates that overall, the AAV-DJ capsid provides a more desirable outcome regardless of the delivery route used.

**Figure 9.**
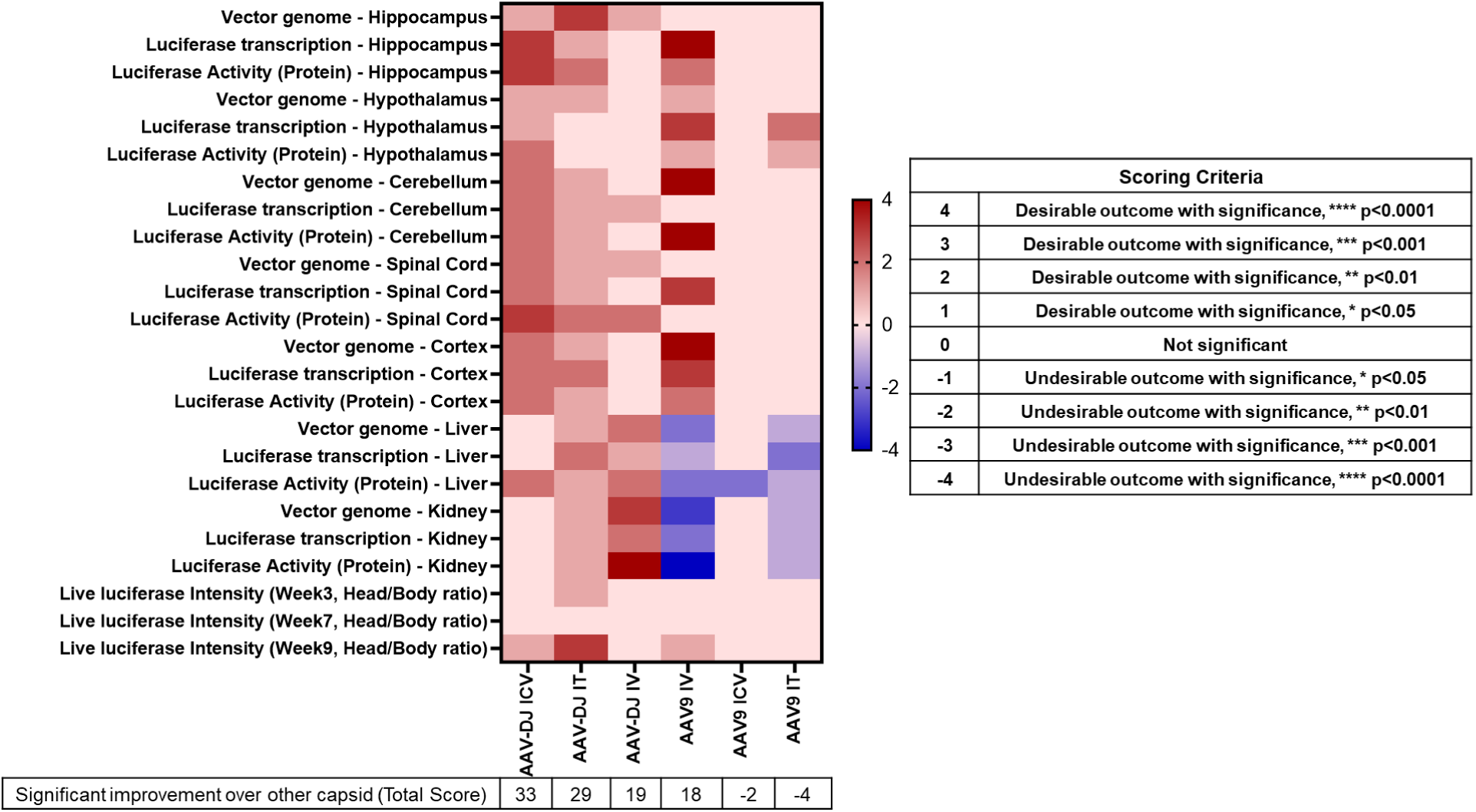
A Heat map summarizing the AAV-mediated gene therapy capsid and delivery route comparison. Scoring criteria are indicated in the right-hand panel based on the statistical significance obtained in each assay as compared to the other capsid.

## Discussion

AAV vectors are being increasingly incorporated into the development of gene therapies for a wide variety of genetically inherited disorders due to their ability to efficiently transduce various cell types, including non-dividing cells with relatively minimal immunogenicity (39–42). AAV9 has demonstrated broad tropism for a variety of organs including the CNS and is the gene delivery vehicle currently being used in the FDA-approved therapy for SMA (7). Despite the overall clinical success with this therapy, there remain challenges due to the occurrence of immunogenicity to natural serotypes such as AAV9. Many investigators are working to develop engineered capsids that evade pre-existing neutralizing antibodies or induce milder immune responses, in the hope that this will allow for more sustained transgene expression and therapeutic benefit (43–45).

The AAV-DJ serotype used in our studies was developed and identified through screening pools of hybrid AAV vectors generated from five starting natural serotypes (AAV 2, 4, 5, 8, and 9) (46). The result, AAV-DJ, is an AAV 2/8/9 chimera that differs from its closest AAV serotype relative (AAV2) by 60 amino acids. Previous studies have shown that AAV-DJ is highly efficient *in vivo* (47–50) and more recently, a non-human primate study demonstrated that AAV-DJ successfully achieved broad transduction throughout the brain(18). A careful assessment of AAV-DJ transduction as compared to AAV9 in mice was needed to provide the rationale for testing this capsid as a gene delivery vehicle to treat neurological disorders in small and large animal models as well as in future human clinical trials.

AAV-DJ exhibits robust transduction efficiency in both *in vitro* and *in vivo* (46), and outperforms several naturally occurring AAV serotypes, including AAV2 and AAV8, with respect to tissue transduction efficiency(46,51,52). The unique transduction profile of AAV-DJ can be attributed to its unique capsid properties, including altered receptor binding and cellular trafficking mechanisms (20,46). Our hypothesis that AAV-DJ would provide more on-target CNS expression and less expression in the liver and kidney turned out to be true when directly compared to results using the same doses and delivery routes for AAV9.

We compared the transduction efficiency of AAV9 and AAV-DJ using three delivery routes (IV, IT, and ICV) to characterize differences and similarities in their abilities to transduce specific regions of the brain, the spinal cord, specific neuronal cell types of interest, and off target organs (liver and kidney). A dose of 1x10^13^ vg/kg was used for IV administrations of these vectors as this aligns well with typical systemic doses used in previous AAV mediated gene therapy studies (28,53). A lower dose of 1x10^12^ vg/kg was used for both IT and ICV injections. Thus, we remained within the typical dose range for these delivery routes due to volume limitations when performing these types of studies in mice (54).

Improved targeting efficiency of neurological gene therapies is crucial because high liver innate and adaptive immune responses are often observed even in successful AAV gene therapies (23,55). Our use of the ubiquitous chicken-β-actin (CBA) promoter to express luciferase-YFP fusion construct enabled us to detect luciferase activity *in vivo* at different time points and enabled us to conduct IF imaging of expression using an antibody that recognized YFP. IVIS bioluminescence imaging enabled us to conduct non-invasive longitudinal assessments of general transgene expression on live animals (56). This methodology is easily accessible and simple to perform to gain a broad and general sense of transgene expression (57). However, high expression levels near the skin in one area can skew results such that, due to detection gradient scaling, the deeper location of expression as in the brain or other areas can be masked in images. As IVIS software can detect this bioluminescence and provide data from selected regions of interest despite masking, the broad-based transgene expression information from these studies is valuable despite its regional nature.

Future longitudinal studies designed to test AAV-DJ in large animal models such as swine, which more accurately match the size and complexity of the human brain, will be necessary to optimize doses and volumes. As promoters also provide a level of control over vector transgene expression, future studies designed to pair the AAV-DJ capsid and optimal delivery route with promoter comparisons will further refine CNS targeted AAV-mediated gene therapy strategies.

In general, RNA transcript and protein expression levels were highly similar to those observed in the vg data, with the few differences likely related to variances in the distribution of data between cohorts. Differences in correlation can arise due to various factors such as post-transcriptional modifications, protein stability, and translational efficiency (58). Additionally, differences in assay sensitivity and specificity have been shown to contribute to variations in data output. Therefore, while these analyses provide insights into gene therapy and molecular biology, researchers can interpret and choose a capsid according to their specific targets and study objectives.

In conclusion, our study indicates that AAV-DJ is superior to AAV9 for targeting of brain and spinal cord while concurrently de-targeting liver and kidney, for all three delivery routes examined in mice. These findings enable the selection of a combined optimal capsid and administration route for various CNS disease-specific therapeutic targets. This study also establishes a platform upon which future studies can be designed to evaluate AAV-DJ capsid modifications for improved neuronal transduction and promoter comparisons to facilitate the efficient and effective delivery of therapeutic transgenes.

## Acknowledgements

This work was supported by the resources and staff at the University of Minnesota University Imaging Centers (UIC: SCR_020997). Thank you to CFI/ITN Imaging Core and Dr. Jason Mitchell for the microscope training and to Alexia Hang and Frances Sehneah for assistance in RNA and gDNA isolation. Figure 1 was created with the help of BioRender.com.

## Author Contributions

M.C. and C.A.P. designed the experiments, analyzed the data, and wrote the manuscript with input and editing from P.B.K., G.A., J.A.T, and A.L.D. Virus production, purification and its quality controls were performed by O.F.D. and G.A. AAV administrations and IVIS imaging were performed by M.C., A.L.D. and F.K., M.C., and A.L.D. performed quantitative PCRs and luciferase assays. M.C. and A.L.D. performed histology preparations and M.C. performed immunofluorescence assays. A.L.D., A.H. quantified the immunofluorescence assays blinded. All authors reviewed a draft of the manuscript and provided input.

---

**Supplementary Figure 1.**
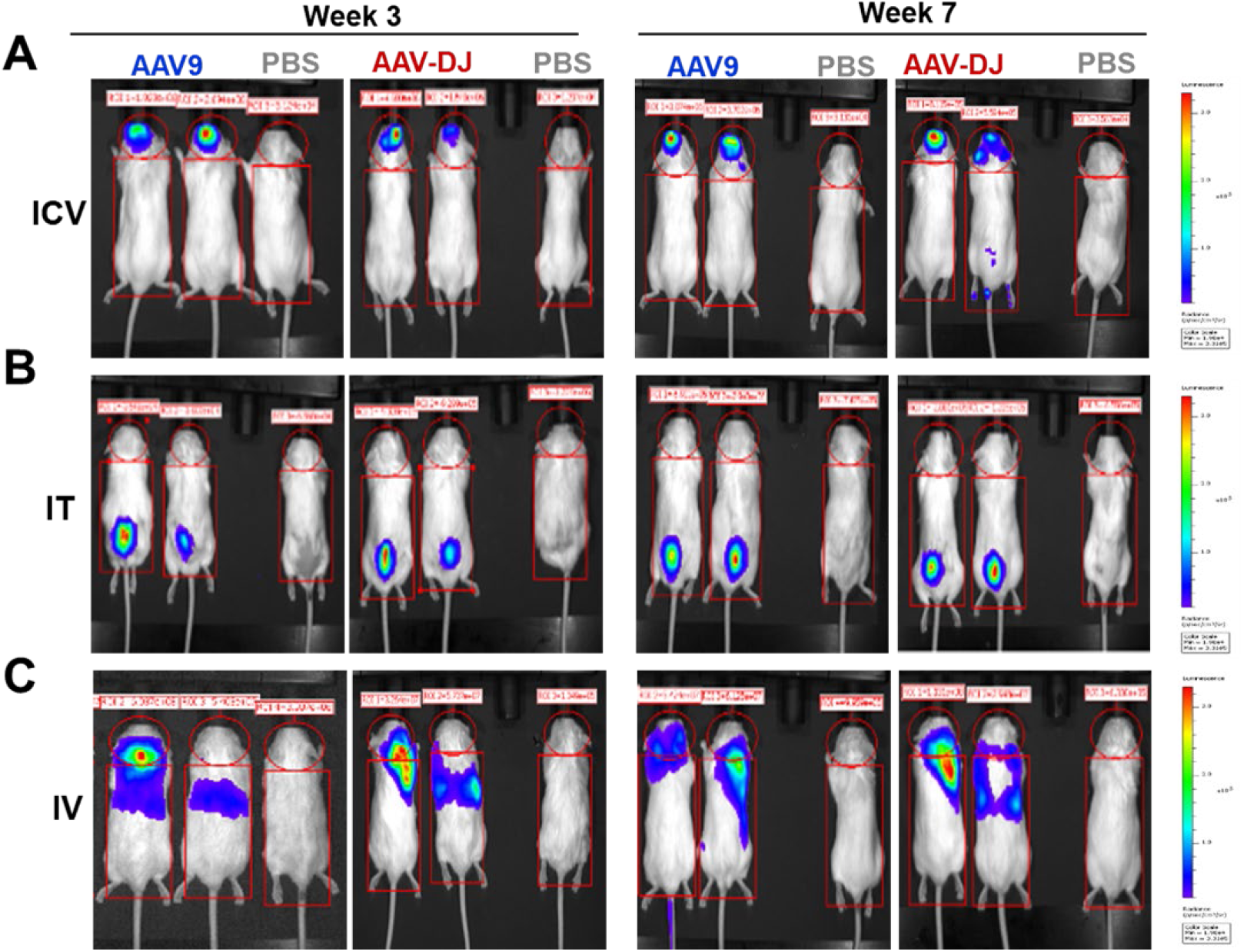
Representative images of two mice from AAV9 and AAV-DJ groups representing each delivery route cohort for week 3 and week 7 (Figure 1). The mouse on the right-most side of every image is a No virus control.

**Supplementary Figure 2.**
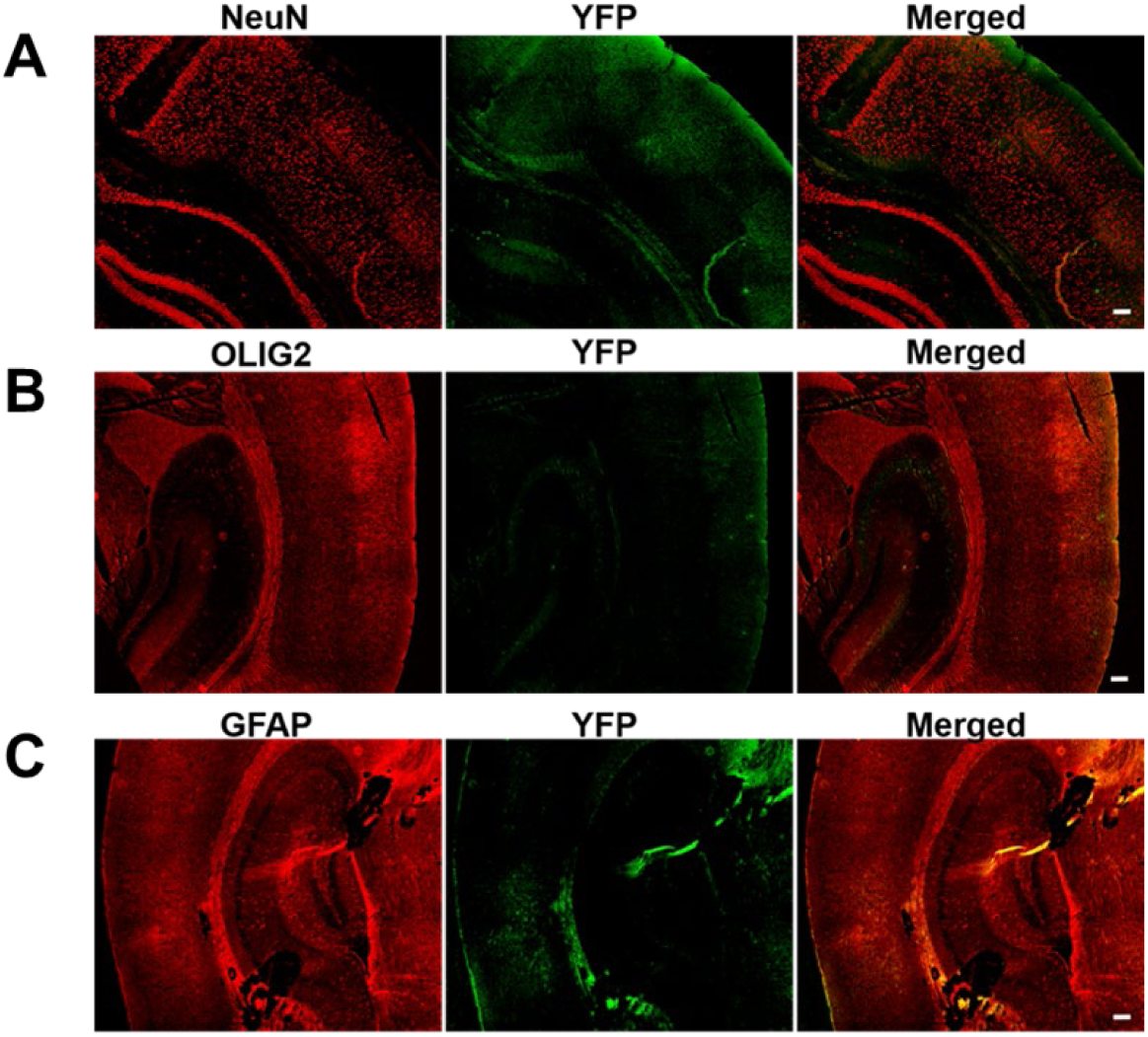
Representative immunofluorescence image of 1xPBS injected mouse brain YFP (green) immunostaining with different markers A. NeuN, B. OLIG2 and C. GFAP (scale bar 100 µm). These images are negative/background controls acquired using the same exposure times and settings as those images from treated mice.

**Supplementary Figure 3.**
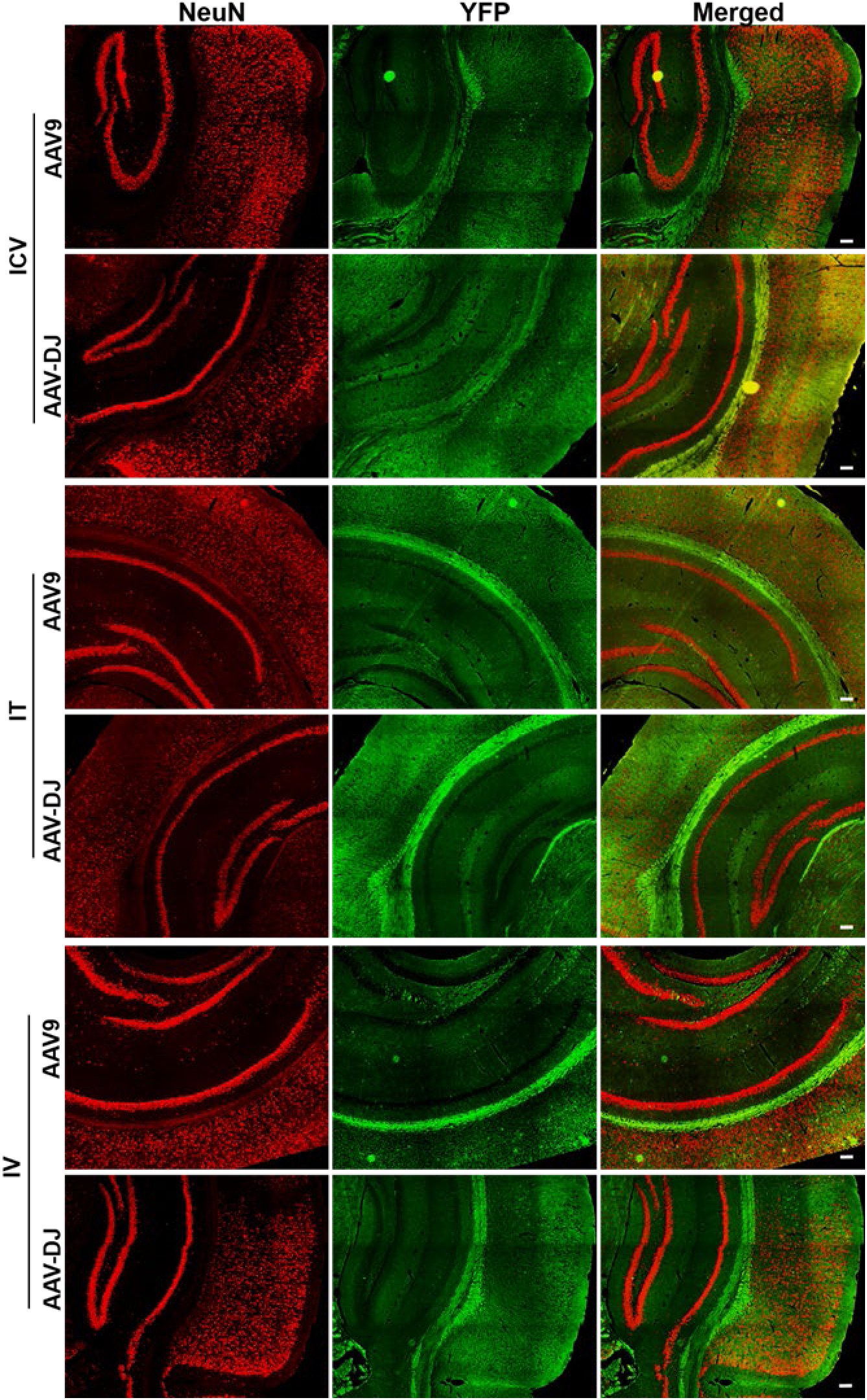
Representative enlarged immunofluorescence images of mouse brains showing NeuN (red), YFP (green) and merged (scale bar 100 µm) (Figure 6).

**Supplementary Figure 4.**
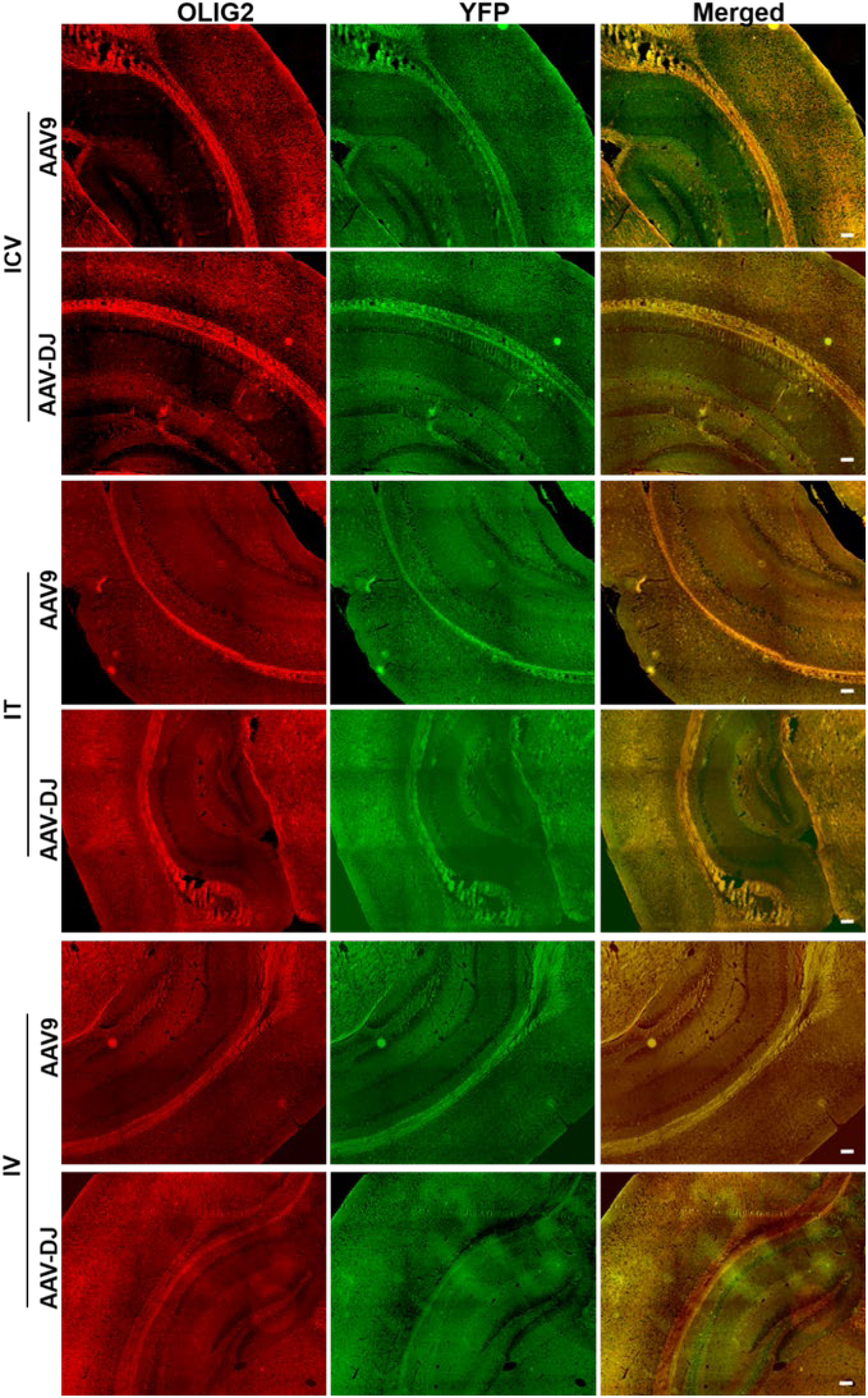
Representative enlarged immunofluorescence images of mouse brains showing OLIG2 (red), YFP (green) and merged (scale bar 100 µm) (Figure 7).

**Supplementary Figure 5.**
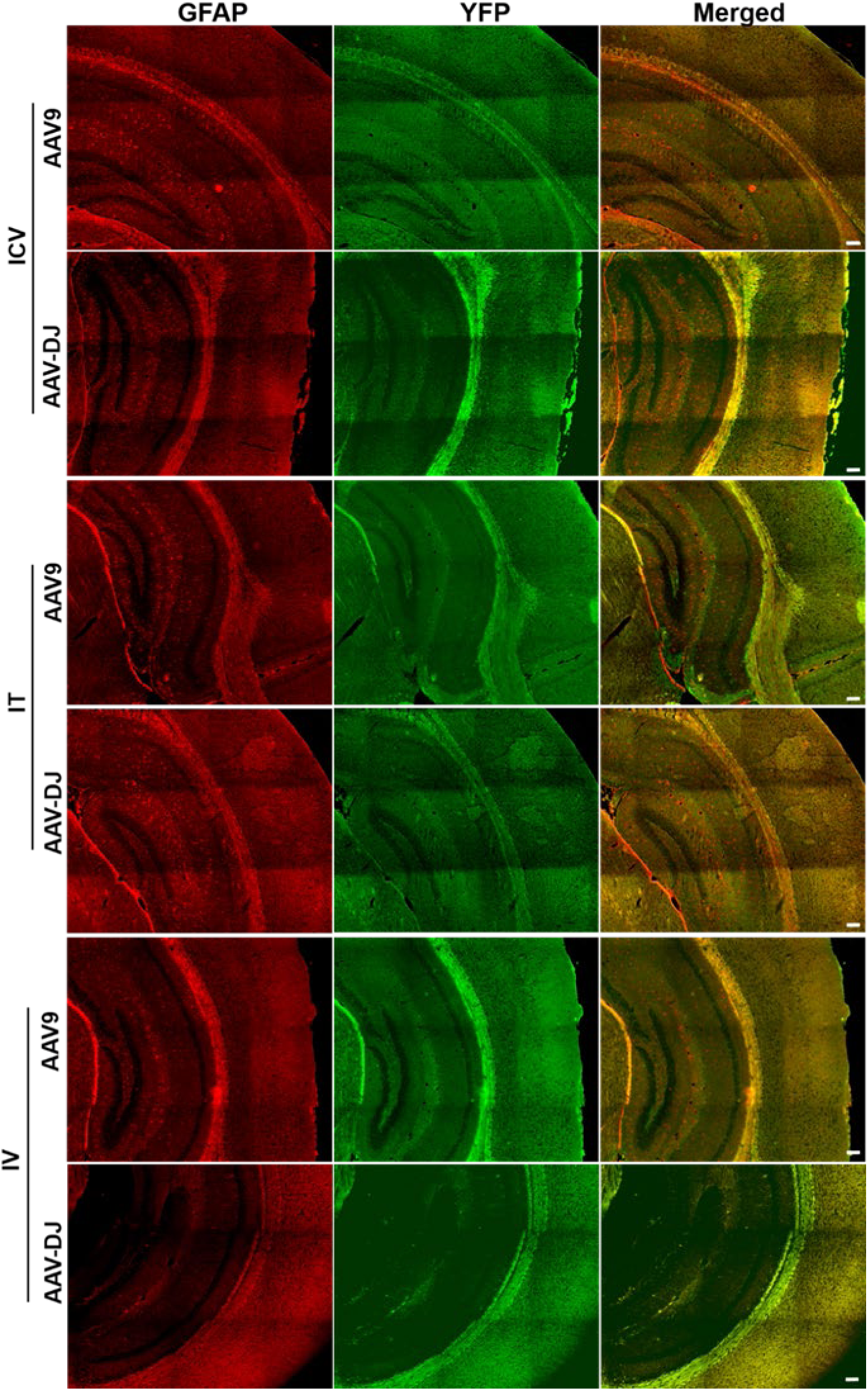
Representative enlarged immunofluorescence images of mouse brains showing GFAP (red), YFP (green) and merged (scale bar 100 µm) (Figure 8).

**Supplementary Figure 6.**
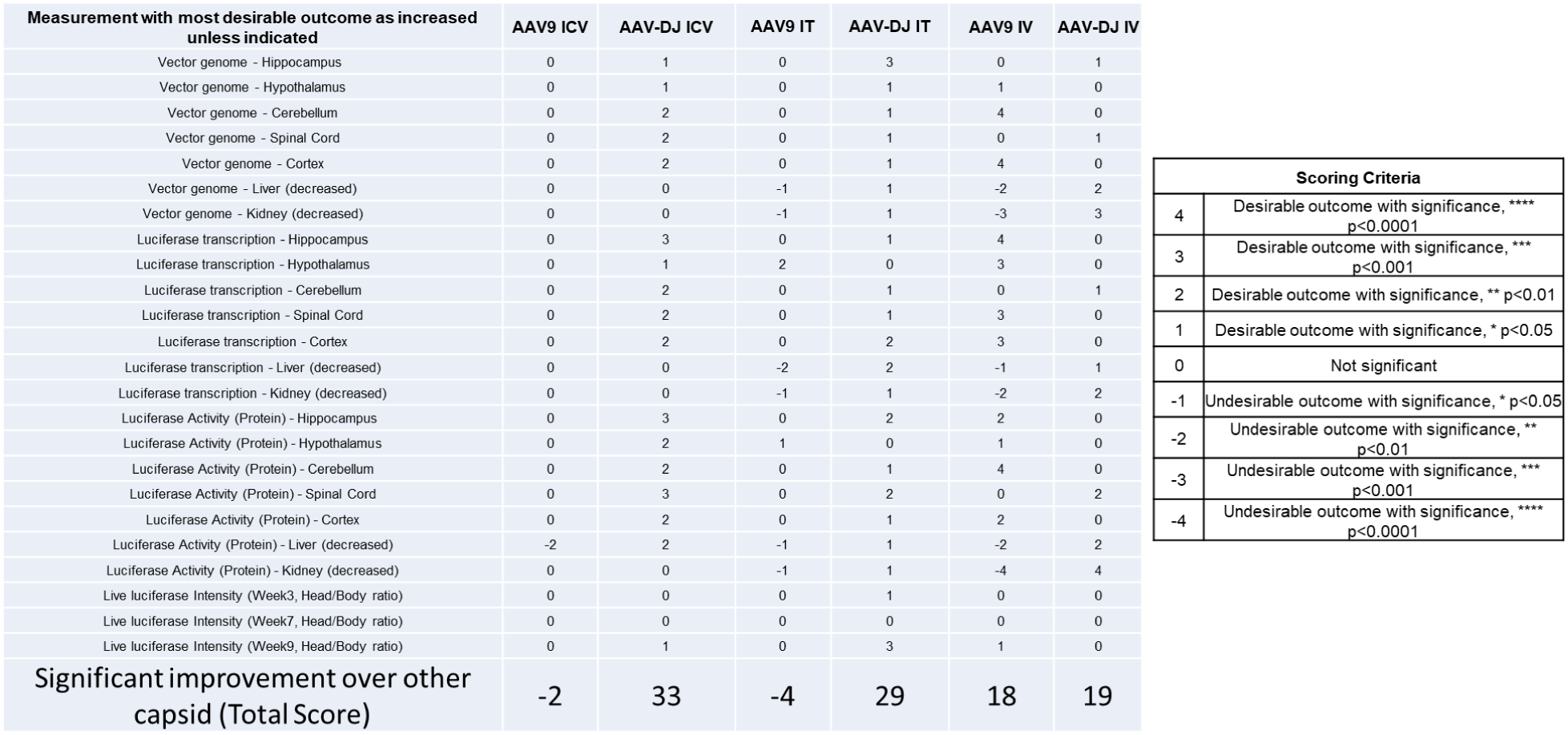
AAV-mediated gene therapy capsid and delivery route comparison summary score table. Scoring criteria is indicated in the right panel.

